# Nrf2 represses ER stress-related ferroptosis in renal carcinoma cells via HO-1

**DOI:** 10.1101/2020.03.04.977769

**Authors:** Yuanqing Shen, Zhiquan Hu, Jihua Tian, Jiahua Gan, Chunjin Ke, Chunguang Yang, Xing Zeng, Guanxin Shen

**Affiliations:** Department of Urology, Tongji Hospital affiliated Tongji Medical College of Huazhong University of Science and Technology (HUST), 1095 Jiefang Avenue, Wuhan, 430030, China; Department of Immunology, Tongji Medical College of Huazhong University of Science and Technology (HUST), No. 13 Hangkong Road, Hankou, Wuhan, 430030, People’s Republic of China

**Keywords:** ferroptosis, endoplasmic reticulum stress, nuclear factor-E2-related factor 2, heme oxygenase-1, renal cell carcinoma

## Abstract

Selective-induction of regulated cell death is considered as a useful strategy for developing effective therapies against renal cell carcinoma (RCC). In this study, the role of nuclear factor-E2-related factor 2 (Nrf2) and its downstream genes in erastin-induced ferroptosis of renal carcinoma cells was investigated. Erastin induced endoplasmic reticulum (ER) stress and ferroptosis in renal carcinoma cells (786-0 and Caki-1), which was confirmed by morphological changes, increased lipid oxidation and ferroptosis inhibition. Data obtained from TCGA dataset showed that Nrf2 and heme oxygenase-1 (HO-1) were not only significantly differentially expressed between RCC and normal samples, but also associated with better survival in RCC patients. Moreover, the process of ferroptosis induced by erastin in renal carcinoma cells was accompanied by increased expression of Nrf2 and HO-1. Inhibition of Nrf2 or HO-1 expression enhanced lipid ROS accumulation and ferroptosis. Furthermore, pharmacological activation of HO-1 reversed ferroptosis-sensitizing effect of Nrf2 silencing, thereby clarifying the key role of Nrf2/HO-1 pathway in erastin-induced ferroptosis in renal carcinoma cells. In summary, we identified the role of Nrf2/HO-1 signaling pathway in the protection of renal carcinoma cells from ER stress-related ferroptosis, which indicated that targeting of Nrf2/HO-1 pathway as a viable treatment strategy for RCC.

Graphical abstract (supplementary figure):
Nrf2 protects against ER stress-related ferroptosis via HO-1 in renal cancer cells. Cystine/glutamate antiporter (SLC7A11) abbreviated as xCT. Mitochondria abbreviated as Mito.

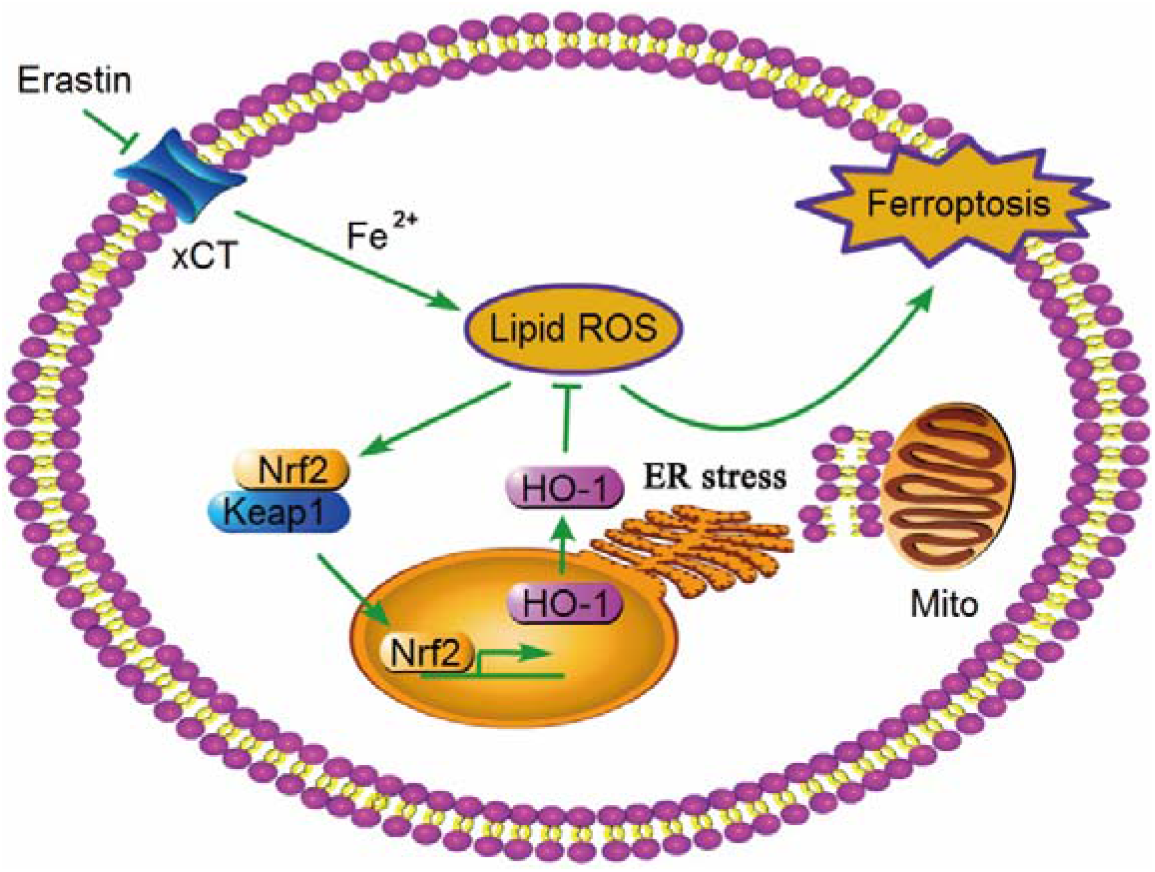

## Introduction

Renal cell carcinoma (RCC) is a common and fetal form of malignancy [1]. Tyrosine kinase inhibitors (TKIs) induces apoptotic cell death, and are approved for treating patients with RCC, but are associated with limited survival benefits [2]. Resistance to current anticancer therapies presents significant unmet medical needs. Induction of selective non-apoptotic cell death might be considered as an effective method for developing more effective therapies against RCC[3].

Ferroptosis is a new form of regulated cell death, and is mediated by accumulation of lipid reactive oxygen species (ROS) in an iron-dependent manner [4]. The other forms of regulated cell death such as apoptosis, necroptosis, and autophagy are morphologically, biochemically, and genetically distinct from ferroptosis. It is characterized by decreased mitochondrial cristae and ruptured outer mitochondrial membrane [5]. RAS mutant tumor cells, including renal carcinoma cells are sensitive to ferroptosis-inducing agents (such as erastin) [6]. Emerging evidence indicated that ferroptosis might be an adaptive process to remove carcinogenic cells [7, 8]. Several modulators of ferroptosis have been linked to certain tumors [9]. Glutathioneperoxidase-4 acts as a key antioxidant in redox regulation, and is a regulator of ferroptosis [10, 11]. p53 promotes ferroptosis by inhibiting cystine/glutamate uptake [8]. Inhibition of cysteine/glutamate exchange by RNA sequencing leads to endoplasmic reticulum (ER) stress and ferroptosis [12]. However, the underlying molecular mechanisms of ferroptosis still remain elusive.

Nuclear factor-E2-related factor 2 (Nrf2) is an essential regulator of cellular oxidative response. Transcriptional activation of its downstream genes plays an important role in maintaining the ROS level [13]. Under normal conditions, Nrf2 is anchored to Kelch-likeECH-associated protein 1 (Keap1) and is rapidly targeted for degradation. ROS degrades Nrf2 inhibitor Keap1, thereby stabilizing and translocating Nrf2 into the nucleus, in which the inducible expression of many antioxidant-responsive elements (ARE) genes, including Heme oxygenase-1 (HO-1), NAD(P)H:quinone oxidoreductase 1 (NQO1), γ-glutamyl-cysteine synthetase (γ-GCS), etc. are regulated. It is not a surprise that Nrf2 protects many kinds of cells from ROS-mediated cell death. Inhibition of Nrf2 sensitizes hepatocellular carcinoma cells to ferroptosis induced by sorafenib, head and neck cancer cells to ferroptosis induced by artesunate, targeting Nrf2 in the regulation of ferroptosis as a new therapeutic strategy for cancer [14, 15]. Biological function and cellular mechanism of Nrf2 axis in ferroptosis of renal carcinoma cells requires further elucidation.

Hence, in this study, we firstly demonstrated Nrf2 represses ER stress-related ferroptosis in renal carcinoma cells via HO-1. Erastin-induced ferroptosis was accompanied by increased levels of lipid oxidation and antioxidants in renal carcinoma cells. Differential expression of Nrf2 and HO-1 was further validated based on the analysis of public dataset. Genetic silencing studies and pharmacological approaches identified the role of Nrf2/HO-1 signaling pathway in the protection of renal carcinoma cells from ER stress-related ferroptosis.

## Materials and Methods

### Cell Culture and Chemicals

RCC cell lines (786-0 and Caki-1) were obtained from ATCC cell bank. These cells were cultured in 1640 medium (Hyclone, USA) supplemented with 10% fetal calf serum (GIBCO, USA) at 37°C in 5% CO_2_ humidified atmosphere. Erastin (SelleckChem, USA) was dissolved in DMSO (Sigma-Aldrich, USA). The below mentioned antibodies were obtained: GRP78 (Proteintech Group, USA), Nrf2 (Proteintech Group, USA), HO-1 (Proteintech Group, USA), NQO1 (proteintech, USA), Keap1 (Proteintech Group, USA), microtubule-associated protein 1 light chain 3 beta (LC3B) (Proteintech Group, USA), caspase 3 (Proteintech Group, USA), β-actin (Proteintech Group, USA), and HRP-conjugated goat anti-rabbit secondary antibody (Proteintech, USA). Other reagents used were as follows: Ferrostatin-1 (Fer-1) (SelleckChem, USA) and Acetylcysteine (NAC) (SelleckChem, USA), deferoxamine (DFO) (Sigma-Aldrich, USA), cobalt protoporphyrin (CoPP) (Sig-ma-Aldrich, USA), HA15 (MedChemExpress, USA).

### Cell Viability Assay

Cytotoxicity was assessed using MTT assay (Sigma-Aldrich, USA). 786-0 and Caki-1 cells were cultured in a 96 well plates and incubated with erastin alone or in combination with other reagents (Fer-1, DFO, NAC or CoPP) for 24 hours (h). The absorbance was then measured at 570nm according to the standard protocols (Beckman, USA). All experiments were performed in triplicate.

### Stable cell lines

786-0 and Caki-1 cells were seeded. For stable Nrf2 or HO-1 knockdown, a lentiviral vector containing small hairpin RNA (shRNA) directed against Nrf2 or HO-1 or control shRNA and transfecting renal carcinoma cells was constructed according to the manufacturer’s instructions (Transomic, Huntsville, AL, USA). The infected cells were then incubated with 2.0□μg/mL puromycin (Sigma-Aldrich) until stable clones were obtained. The target sequence of shRNA-Nrf2 was 5′-CCGGCCGGCATTTCACTAAACACAACTCGAGTTGTGTTTAGTGAAATGCCGGTTTTT-3′. The target sequence of shRNA-HO-1 was 5′-GATCCGAGAACCACGGTCTGCACCATATTATTCAAGAGATAATATGGTGCAGACCGTGGTTCTCTTTTTTC-3′. Western blotting was performed to confirm shRNA-induced gene silencing.

### Lipid Peroxidation measurement

786-0 and Caki-1cells were seeded for 24 h before treatment at a density of 10^6^ cells per well. After incubation with erastin or other reagents for 24□h, renal carcinoma cells were then collected. Lipid peroxidation (MDA) in the supernatant of renal carcinoma cells lysates was measured by Beckman flow cytometer after adding 2 μM of C11-BODIPY C11 (Invitrogen, USA) for 30 min.

### Western Blotting

Western blotting was performed to analyze protein expression [16]. The following primary antibodies were used: GRP78, Keap1, Nrf2, HO-1, NQO1, LC3B, caspase 3, and β-actin. All antibodies were diluted between 1:500 and 1:1000.

### Scanning Electron Microscopy

The procedures of scanning electron microscopy for ultrastructural inspection are done as described previously [16].

### Fluorescence Microscopy

Incubation procedures for labeling the cells with C11-BODIPY^581/591^ (Lipid Peroxidation Sensor) were performed as described [17]. Nrf2 or HO-1 knockdown in renal carcinoma cells were incubated with 10 μmol/L erastin for 24 h. The samples were examined at 100x magnification.

### Public transcriptome data

Analysis of Nrf2 expression and its transcriptional activity (HO-1, NQO-1, γ-GCS) in RCC was performed using the TCGA (https://tcga-data.nci.nih.gov/tcga) databases for cancer expression data. GCLC and GCLM are catalytic subunits and modifier subunits of γ-GCS, respectively. The expression and survival data from 540 RCC and 72 normal renal tissue samples were used.

### Statistical Analysis

All statistical analysis was performed using SPSS 12.0. Data were presented as means ± SD. One-way analysis of variance (ANOVA) followed by student’s t-test were used to determine statistically significant differences between the test samples and controls. A p-value of less than 0.05 was considered to be statistically significant.

## Results

### Erastin induces ER stress-related ferroptosis in renal carcinoma cells

Erastin decreased the viability of renal carcinoma cells in a dose-dependent manner (Fig. 1A, B), and this effect was blocked by pretreatment with the ferroptosis inhibitor ferrostatin-1 (Fer-1), iron chelator deferoxamine (DFO) or ROS inhibitor acetylcysteine (NAC) (Fig. 1B, D). Moreover, electron microscopic study of renal carcinoma cells demonstrated decreased mitochondrial cristae or ruptured outer mitochondrial membrane after incubation with erastin for 24 h (Fig. 1C). Oxidative stress is considered as a major inducer of Nrf2 expression. Lipid ROS (MDA) levels were increased in renal carcinoma cells treated with erastin (Fig. 1E). The cellular vacuolization and enhanced expression of glucose-regulated protein 78 (GRP78) induced by erastin were reversed by co-incubation of lipid peroxidation inhibitor ferrostatin-1, showing that ER stress was stimulated by lipid ROS (Fig. 1B, F). HA15 is an effective inhibitor of the essential ER chaperone GRP78/BiP [18]. Cytotoxicity of erastin was markedly inhibited by co-incubation of HA15, indicating that ER stress protected renal carcinoma cells from ferroptosis induced by excessive lipid ROS (Fig. 1G). Meanwhile, expression of caspase-3 and microtubule-associated protein 1 light chain 3 beta (LC3B) showed no significant difference in renal carcinoma cells treated with erastin, indicating no activation of apoptosis or autophagy (Fig. 1H). These findings suggested that erastin induced ER stress-related ferroptosis in renal carcinoma cells.

**Figure 1.**
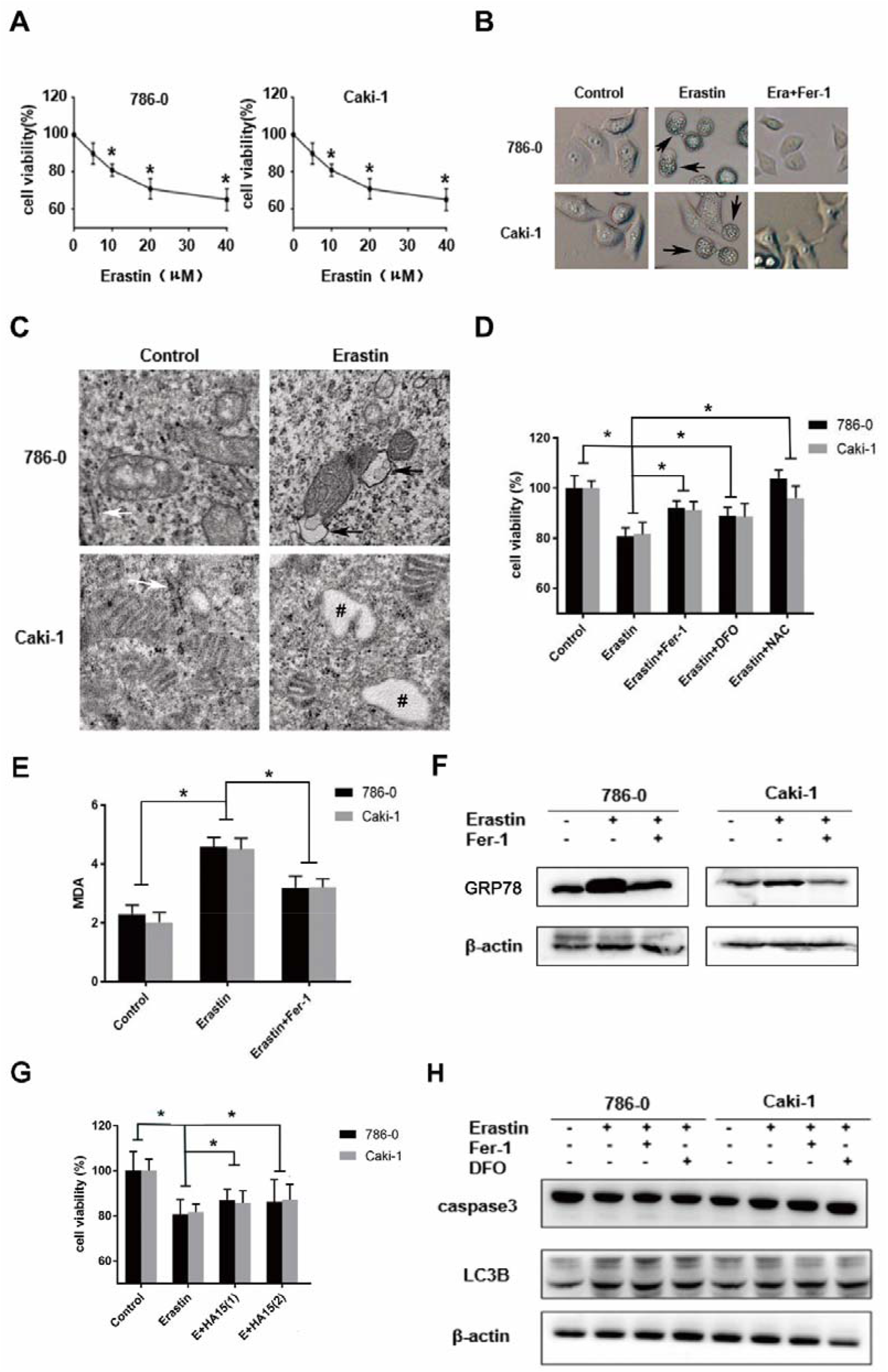
Erastin induces ER stress-related ferroptosis in renal carcinoma cells. (A) Cell viability was assessed by MTT assay after incubation at different concentrations of erastin for 24 h in renal carcinoma cells. (B) Microphotographs showing cellular vacuolization (black arrows) after exposure to 10 μM erastin (Era) with or without 10 μM ferrostatin-1 (Fer-1) for 24 h. (C) Ultrastructural inspection of erastin-treated renal carcinoma cells. Outer membrane rupture of mitochondria is highlighted by black arrows. White arrows point to normal ER. Pounds indicate dilated ER cavity. (D) Renal carcinoma cells were incubated with erastin (10 μM) for 24 h with or without ferrostatin-1 (Fer-1, 10 μM), deferoxamine (DFO, 100 mM) or acetylcysteine (NAC, 10 mM) pretreatment and cell viability was assayed. (E) Elevation of lipid ROS (MDA) in renal carcinoma cells incubated with 10 μM erastin for 24 h. (F) The expression of glucose-regulated protein 78 (GRP78) was analyzed by western blot in renal carcinoma cells after incubated with 10 μM erastin for 24 h with β-actin as an internal control. (G) Renal carcinoma cells were exposed to erastin (E, 10 μM) for 24 h with or without HA15 (2.5 μM and 5 μM, respectively) pretreatment, then cell viability was assayed. (H) Western blot analyses of renal carcinoma cells is incubated with 10 μM erastin for 24 h. Error bars indicate standard error from three technical replicates. * P < 0.05.

### Increased Nrf2/HO-1 expression during ferroptosis correlates with better survival in RCC patients

Differentially expressed genes between RCC and normal samples provided an opportunity to study the regulation of ferroptosis. The expression data from The Cancer Genome Atlas (TCGA) was used to differentially express Nrf2 and its downstream genes. GCLC and GCLM are catalytic subunits and modifier subunits of γ-GCS, respectively. This analysis revealed that Nrf2 and NQO1 were significantly down-regulated in RCC, whereas HO-1 was up-regulated (Fig. 2A). Western blotting analysis revealed significantly increased Nrf2/HO-1 protein level and decreased Keap1 protein level, but not NQO1 in erastin-treated renal carcinoma cells (Fig. 2B), indicating the stabilization of Nrf2 protein in ferroptosis by Keap1 degradation. Moreover, survival analysis of TCGA dataset showed that increased expression of Nrf2 and HO-1 was correlated with better survival in patients with RCC (Fig. 2C). In brief, our findings showed that Nrf2 and HO-1 could play an essential role in ferroptosis regulation in erastin-treated renal carcinoma cells.

**Figure 2.**
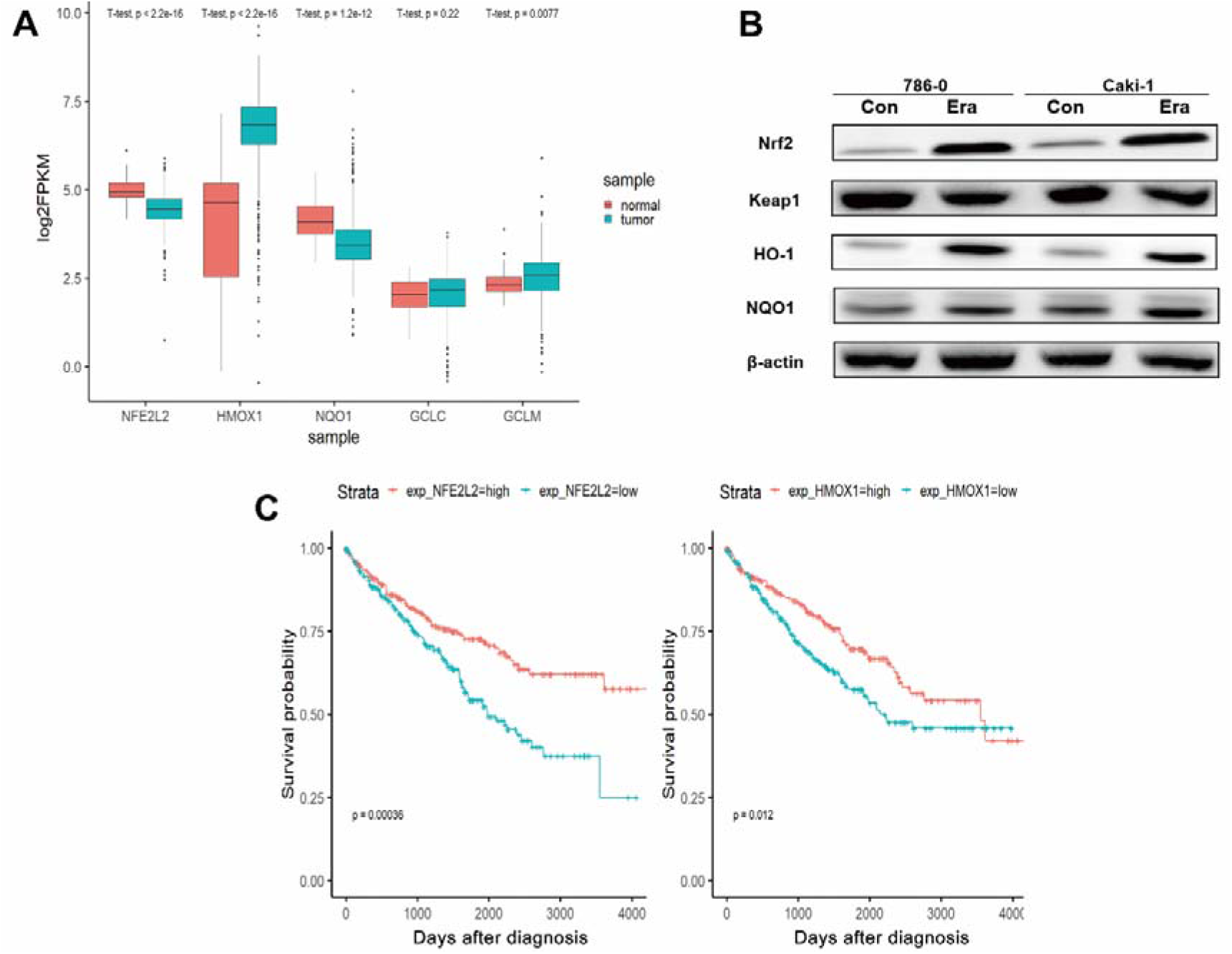
Increased Nrf2/HO-1 expression during ferroptosis correlates with better survival in RCC patients. (A) Analysis of Nrf2 expression and its downstream genes from The Cancer Genome Atlas (TCGA) dataset shows Nrf2 (NFE2L2), HO-1 (HMOX1) and NQO1 are significantly differentially expressed genes between RCC and normal samples (P□<□0.001). (B) Expression of Nrf2, HO-1, NQO1 and Keap1 in renal carcinoma cells treated with 10 μM erastin (Era) for 24 h with β-actin as an internal control (Con). (C) Survival analysis of TCGA dataset shows increased expression of Nrf2 or HO-1 correlates with better survival rate in patients with RCC (P□<□0.05).

### Inhibition of Nrf2 enhances ferroptosis induced by erastin in renal carcinoma cells

Nrf2 gene silencing by specific shRNA significantly decreased the expression of Nrf2 and HO-1 in renal carcinoma cells (Fig. 3D), indicating HO-1 is a downstream target of Nrf2. Cytotoxicity of erastin was markedly enhanced in silenced renal carcinoma cells when compared to shRNA control group (Fig. 3A). The lipid ROS levels were significantly increased in renal carcinoma cells transfected with shNrf2 (Fig. 3B, C). Therefore, inhibition of Nrf2 enhances ferroptosis induced by erastin in renal carcinoma cells.

**Figure 3.**
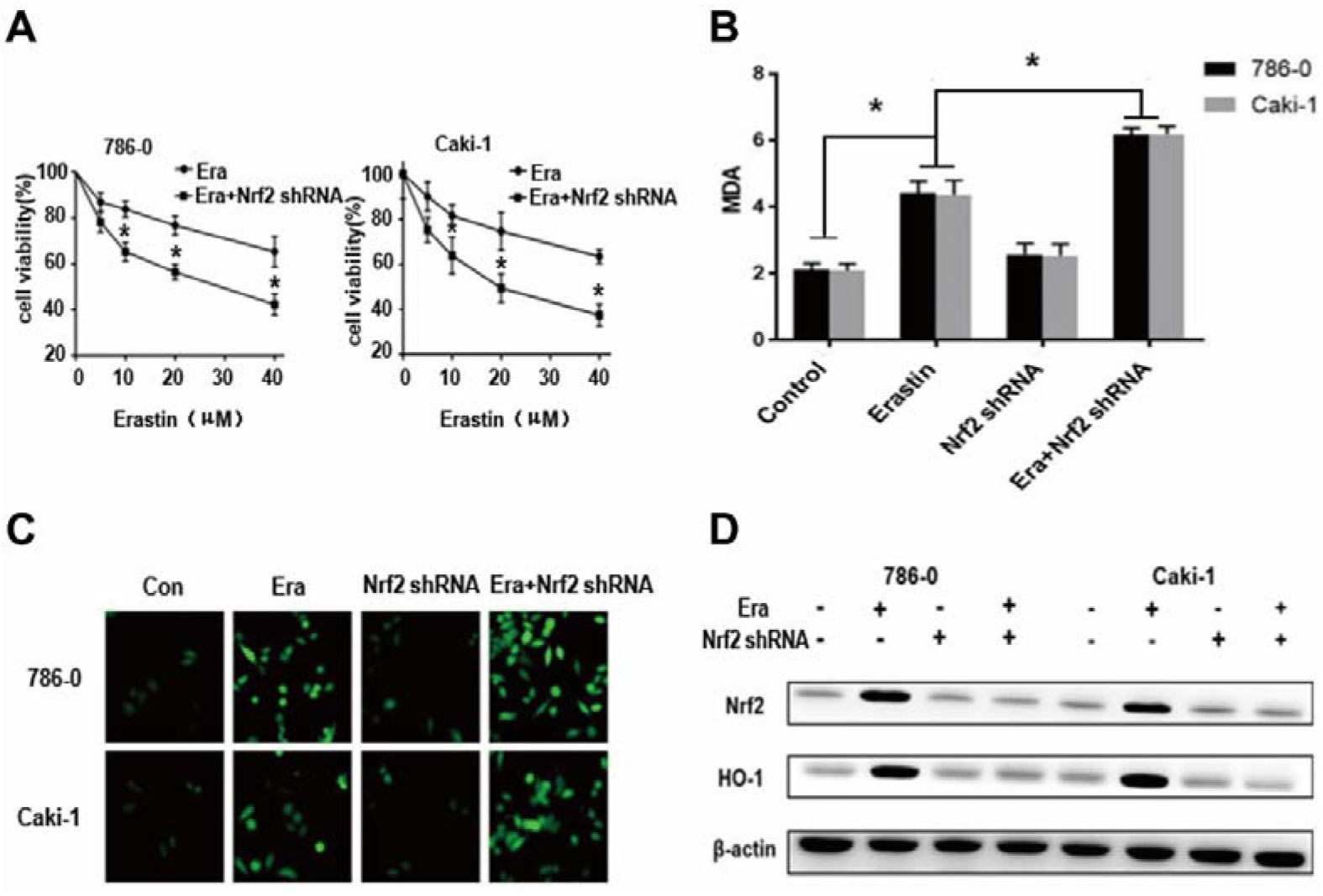
Inhibition of Nrf2 sensitizes renal carcinoma cells to ferroptosis induced by erastin. (A) Nrf2 knocked down renal carcinoma cells treated with erastin (Era, 10 μM) for 24 h and assessed by MTT. (B) Lipid ROS (MDA) was elevated in Nrf2 knockdown of renal carcinoma cells when exposed to 10 μM erastin for 24 h. (C) Nrf2 knockdown of renal carcinoma cells were treated with erastin (10 μM) for 24 h and labeled with C11-BODIPY^581/591^ for detecting lipid peroxidation processes in membranes. (D) Western blot analyses of Nrf2 knockdown in renal carcinoma cells incubated with erastin for 24 h with β-actin as an internal control. Error bars indicate standard error from three technical replicates. * P < 0.05.

### Inhibition of HO-1 sensitizes renal carcinoma cells to ferroptosis induced by erastin

Silencing of HO-1 genes by specific shRNA decreased the expression of HO-1 but not Nrf2 (Fig. 4D). Meanwhile, the cell viability (Fig. 4A) of renal carcinoma cells was more significantly decreased in shHO-1 group than shRNA control group after exposure to erastin. The lipid ROS levels were significantly increased in renal carcinoma cells transfected with shHO-1 (Fig. 4B, C). Therefore, HO-1 expression contributes to the resistance to erastin induced ferroptosis.

**Figure 4.**
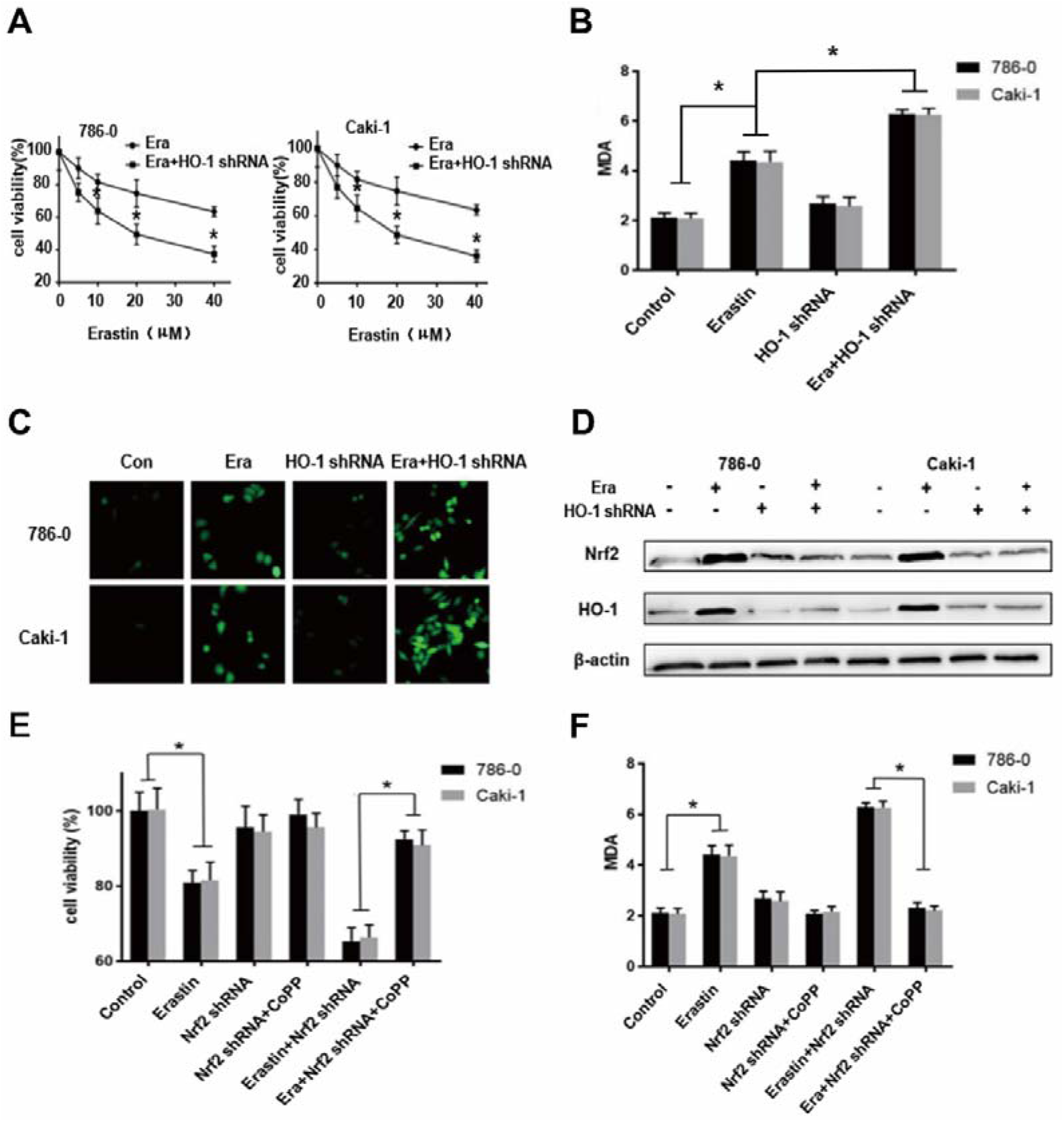
Inhibition or activation of HO-1 determines the sensitivity of renal carcinoma cells to erastin. (A) HO-1 knockdown renal carcinoma cells treated with erastin (Era, 10 μM) for 24 h and assessed by MTT. (B) Lipid ROS (MDA) was elevated in HO-1 knockdown of renal carcinoma cells when exposed to 10 μM erastin for 24 h. (C) HO-1 knockdown of renal carcinoma cells were treated with erastin (10 μM) for 24 h and labeled with C11-BODIPY^581/591^ for lipid peroxidation detection. (D) Western blot analyses of HO-1 knockdown in renal carcinoma cells exposed to erastin for 24 h with β-actin as an internal control. (E) Nrf2 knocked down renal carcinoma cells were treated with erastin (10 μM) for 24 h with or without pretreatment with HO-1 inducer cobalt protoporphyrin (CoPP, 10 μM) and cell viability were assayed. (F) Nrf2 knocked down renal carcinoma cells were treated with erastin (10 μM) for 24 h with or without pretreatment with HO-1 inducer cobalt protoporphyrin (CoPP, 10 μM) and lipid ROS (MDA) was assayed. Error bars indicate standard error from three technical replicates. * P < 0.05.

### HO-1 activation reversed ferroptosis-sensitizing effect of Nrf2 silencing

To further investigate the role of Nrf2/HO-1 signaling pathway in ferroptosis, we explore HO-1 activation in renal carcinoma cells during ferroptosis. Activation of HO-1 by cobalt protoporphyrin (CoPP) significantly reversed erastin-induced and Nrf2 silencing sensitized cell viability decrement (Fig. 4E) and lipid ROS production (Fig. 4F). This suggested that Nrf2 protects erastin-induced ferroptosis in renal carcinoma cells mainly through the induction of HO-1.

## Discussion

Ferroptosis provides a new approach for developing novel targeted therapy. Sensitizing of ferroptosis-inducing agent is considered as a promising strategy for overcoming drug resistance[19]. This study demonstrated that erastin-induced ferroptosis in renal carcinoma cells is accompanied by an increase in lipid oxidation and antioxidants. Genetic inhibition of Nrf2 or HO-1 enhanced ROS accumulation and ferroptosis when co-administered with erastin in renal carcinoma cells. Pharmacological activation of HO-1 identified Nrf2/HO-1 signaling pathway in erastin-induced ferroptosis.

Accumulation of lipid ROS is regarded as a driving factor for ferroptosis [4, 20]. Nrf2 is the central regulator of cellular oxidative response, Nrf2 inhibition has been shown to sensitize only a limited number of cancer cells to ferroptosis [14, 15]. The expression data from TCGA dataset shows Nrf2 is significantly differentially expressed and associated with survival in patients with RCC (Fig 2). The current study, not surprisingly, showed that erastin treatment induced increased lipid ROS accumulation and Nrf2 expression during ferroptosis in renal carcinoma cells (Fig 1E and 2B). Similar to the previous reports [14, 15], genetic silencing of Nrf2 significantly increased lipid ROS levels and sensitized renal carcinoma cells when treated with erastin (Fig 3). These results indicated that the balance between lipid ROS and anti-oxidant remains crucial to ferroptosis. Thus, targeting Nrf2 modulates lipid peroxidation and ferroptosis is a possible strategy for RCC intervention.

Nrf2 defenses against oxidative stress depending on the inducible expression of many ARE genes (HO-1, NQO1, γ-GCS, etc). Expression data from TCGA has shown that both HO-1 and NQO1 are differentially expressed between RCC and normal samples (Fig 2A). Increased expression of HO-1 was correlated with better survival in patients with RCC (Fig 2C). Moreover, HO-1 protein levels were significantly increased in erastin-treated renal carcinoma cells, but not NQO1 (Fig. 2B). This indicated that HO-1 is a key downstream effector molecule of Nrf2-ARE signaling pathway in erastin-treated renal carcinoma cells. HO-1 is a well-known target gene regulated by Nrf2 [21]. It showed a cytoprotective effect against oxidative response, whereas its function in ferroptosis is still not very clear [22]. Upregulation of HO-1 induced cellular ferrous accumulation and ferroptosis in breast cancer cell lines by BAY11-7085 [23]. In contrast, HO-1 mitigates ferroptosis in hepatocellular carcinoma cells and renal proximal tubule cells. This indicated that the function of HO-1 in ferroptosis depends on the inducer and cell type [14, 24]. Genetic silencing of HO-1 showed an increase in lipid ROS levels and sensitized renal carcinoma cells to erastin (Fig 4A-D), suggesting cytoprotective effect of HO-1 in renal carcinoma cells. Furthermore, HO-1 activation reversed the sensitizing effect of Nrf2 silencing on ferroptosis (Fig 4E, F), which demonstrated that the antiferroptotic activity of Nrf2 in renal carcinoma cells mainly depends on the induction of HO-1 as antioxidants. The exact mechanism by which HO-1 suppresses ferroptosis remains to be further studied.

Regulated cell deaths are frequently associated with oxidative injury and ER stress [25]. Prolonged ER stress has been shown to engage a distinct set of pro-apoptotic outputs [26]. Autophagy can be activated by sorafenib induced ER stress through the lysosomal pathway [27]. Moreover, studies have shown that ER stress-mediated Nrf2 activation inhibit inflammation caused by excessive ROS [28]. In our study, ER stress induced by erastin was reversed by co-incubation of ferrostatin-1 (Fig. 1B, F), showing that ER stress was stimulated by lipid ROS. Moreover, ferroptosis inhibition by HA15 indicated that ER stress protected renal carcinoma cells from ferroptosis induced by excessive lipid ROS (Fig. 1G). But the exact role of ROS-dependent ER stress in Nrf2 activation remains to be explored.

In summary, we demonstrated that Nrf2 protects against ER stress-related ferroptosis via HO-1 in renal carcinoma cells. Firstly, we demonstrated that erastin-induced ferroptosis is accompanied by increased lipid oxidation and lipid oxidation-related protective ER stress in renal carcinoma cells. Secondly, genetic inhibition of Nrf2 or HO-1 enhanced lipid peroxidation and ferroptosis when co-administered with erastin in lipid oxidation and lipid oxidation-related ER stress. Furthermore, pharmacological activation of HO-1 identified the key role of Nrf2/HO-1 signaling pathway in erastin-induced ferroptosis. Functional identification of Nrf2/HO-1 signaling pathway in ferroptosis might provide a viable treatment strategy for RCC.

## Acknowledgments

Not applicable.

## Availability of data and materials

The datasets used and/or analyzed during the present study are available from the corresponding author upon reasonable request

## Authors’ Contributions

Conceptualization, writing original draft, Chunguang Yang and Yuanqing Shen;

Date curation, Yuanqing Shen, Jihua Tian, Jiahua Gan, and Chunjin Ke;

Project administration, Zhiquan Hu and Guanxin Shen;

Formal analysis, Xing Zeng;

Writing-review & editing, Zhiquan Hu;

Funding acquisition, Chunguang Yang;

All authors read and approved the final manuscript.

## Ethics approval and consent to participate

Not applicable.

## Patient consent for publication

Not applicable.

## References

1. Bray, F., et al., Global cancer statistics 2018: GLOBOCAN estimates of incidence and mortality worldwide for 36 cancers in 185 countries. CA Cancer J Clin, 2018. 68(6): p. 394–424.

2. Porta, C., et al., The adjuvant treatment of kidney cancer: a multidisciplinary outlook. Nat Rev Nephrol, 2019. 15(7): p. 423–433.

3. Martin-Sanchez, D., et al., Cell death-based approaches in treatment of the urinary tract-associated diseases: a fight for survival in the killing fields. Cell Death Dis, 2018. 9(2): p. 118.

4. Dixon, S.J., et al., Ferroptosis: an iron-dependent form of nonapoptotic cell death. Cell, 2012. 149(5): p. 1060–72.

5. Gao, M., et al., Role of Mitochondria in Ferroptosis. Mol Cell, 2019. 73(2): p. 354–363 e3.

6. Shimada, K., et al., Cell-Line Selectivity Improves the Predictive Power of Pharmacogenomic Analyses and Helps Identify NADPH as Biomarker for Ferroptosis Sensitivity. Cell Chem Biol, 2016. 23(2): p. 225–235.

7. Dixon, S.J., Ferroptosis: bug or feature? Immunol Rev, 2017. 277(1): p. 150–157.

8. Jiang, L., et al., Ferroptosis as a p53-mediated activity during tumour suppression. Nature, 2015. 520(7545): p. 57–62.

9. Mou, Y., et al., Ferroptosis, a new form of cell death: opportunities and challenges in cancer. J Hematol Oncol, 2019. 12(1): p. 34.

10. Yang, W.S., et al., Regulation of ferroptotic cancer cell death by GPX4. Cell, 2014. 156(1–2): p. 317–331.

11. Zou, Y., et al., A GPX4-dependent cancer cell state underlies the clear-cell morphology and confers sensitivity to ferroptosis. Nat Commun, 2019. 10(1): p. 1617.

12. Dixon, S.J., et al., Pharmacological inhibition of cystine-glutamate exchange induces endoplasmic reticulum stress and ferroptosis. Elife, 2014. 3: p. e02523.

13. Rojo de la Vega, M., E. Chapman, and D.D. Zhang, NRF2 and the Hallmarks of Cancer. Cancer Cell, 2018. 34(1): p. 21–43.

14. Sun, X., et al., Activation of the p62-Keap1-NRF2 pathway protects against ferroptosis in hepatocellular carcinoma cells. Hepatology, 2016. 63(1): p. 173–84.

15. Roh, J.L., et al., Nrf2 inhibition reverses the resistance of cisplatin-resistant head and neck cancer cells to artesunate-induced ferroptosis. Redox Biol, 2017. 11: p. 254–262.

16. Yang, C., et al., Curcumin induces apoptosis and protective autophagy in castration-resistant prostate cancer cells through iron chelation. Drug Des Devel Ther, 2017. 11: p. 431–439.

17. Drummen, G.P., et al., C11-BODIPY(581/591), an oxidation-sensitive fluorescent lipid peroxidation probe: (micro)spectroscopic characterization and validation of methodology. Free Radic Biol Med, 2002. 33(4): p. 473–90.

18. Ronco, C., et al., Structure activity relationship and optimization of N-(3-(2-aminothiazol-4-yl)aryl)benzenesulfonamides as anti-cancer compounds against sensitive and resistant cells. Bioorg Med Chem Lett, 2017. 27(10): p. 2192–2196.

19. Friedmann Angeli, J.P., D.V. Krysko, and M. Conrad, Ferroptosis at the crossroads of cancer-acquired drug resistance and immune evasion. Nat Rev Cancer, 2019. 19(7): p. 405–414.

20. Stockwell, B.R., et al., Ferroptosis: A Regulated Cell Death Nexus Linking Metabolism, Redox Biology, and Disease. Cell, 2017. 171(2): p. 273–285.

21. Furfaro, A.L., et al., The Nrf2/HO-1 Axis in Cancer Cell Growth and Chemoresistance. Oxid Med Cell Longev, 2016. 2016: p. 1958174.

22. Chiang, S.K., S.E. Chen, and L.C. Chang, A Dual Role of Heme Oxygenase-1 in Cancer Cells. Int J Mol Sci, 2018. 20(1).

23. Chang, L.C., et al., Heme oxygenase-1 mediates BAY 11-7085 induced ferroptosis. Cancer Lett, 2018. 416: p. 124–137.

24. Adedoyin, O., et al., Heme oxygenase-1 mitigates ferroptosis in renal proximal tubule cells. Am J Physiol Renal Physiol, 2018. 314(5): p. F702-F714.

25. Zhang, Z., et al., Redox signaling and unfolded protein response coordinate cell fate decisions under ER stress. Redox Biol, 2018.

26. Chang, T.K., et al., Coordination between Two Branches of the Unfolded Protein Response Determines Apoptotic Cell Fate. Mol Cell, 2018. 71(4): p. 629–636 e5.

27. Shi, Y.H., et al., Targeting autophagy enhances sorafenib lethality for hepatocellular carcinoma via ER stress-related apoptosis. Autophagy, 2011. 7(10): p. 1159–72.

28. Keleku-Lukwete, N., et al., Amelioration of inflammation and tissue damage in sickle cell model mice by Nrf2 activation. Proc Natl Acad Sci U S A, 2015. 112(39): p. 12169–74.

